# Aberrant Fucosylation of Saliva Glycoprotein Defining Lung Adenocarcinomas Malignancy

**DOI:** 10.1101/2021.12.16.472888

**Authors:** Ziyuan Gao, Zhen Wu, Ying Han, Xumin Zhang, Piliang Hao, Mingming Xu, Shan Huang, Shuwei Li, Jun Xia, Junhong Jiang, Shuang Yang

## Abstract

Aberrant glycosylation is a hallmark of cancer found during tumorigenesis and tumor progression. Lung cancer induced by oncogene mutations has been detected in the patient’s saliva, and saliva glycosylation has been altered. Saliva contains highly glycosylated glycoproteins, the characteristics of which may be related to various diseases. Therefore, elucidating cancer-specific glycosylation in the saliva of healthy, non-cancer, and cancer patients can reveal whether tumor glycosylation has unique characteristics for early diagnosis. In this work, we used a solid-phase chemoenzymatic method to study the glycosylation of saliva glycoproteins in clinical specimens. The results showed that the α1,6-core fucosylation of glycoproteins in cancer patients was significant increased. The fucosylation of α1,2 or α1,3 is also increased in cancer patients. We further analyzed the expression of fucosyltransferases responsible for α1,2, α1,3, α1,6 fucosylation. The fucosylation of the saliva of cancer patients is drastically different from that of non-cancer or health controls. These results indicate that the glycoform of saliva fucosylation distinguishes lung cancer from other diseases, and this feature has the potential to diagnose lung adenocarcinoma.

**TOC:** Fucosylation biosynthesis in lung cancer. Saliva fucosylation contains α1,2-linked, α1,3-linked, α1,6-linked fucosylation in lung cancer.

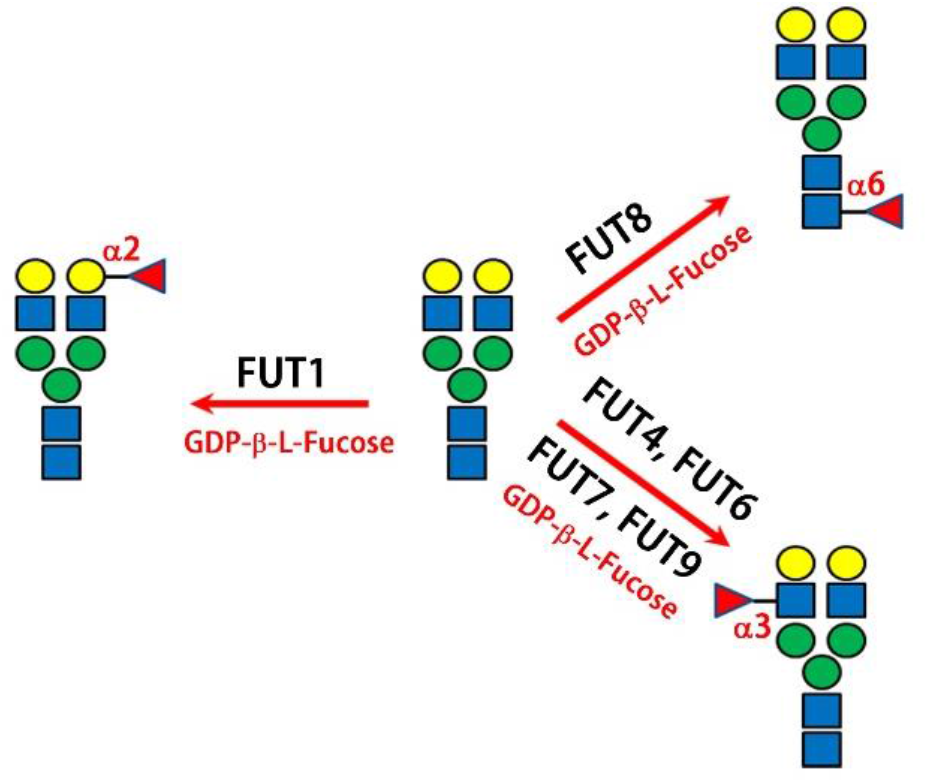

## Introduction

As one of the common post-translationally modifications, glycosylation is associated with many diseases, and its abnormal changes can affect the pathophysiology of cells or organisms (Rudd *et al*, 2001; Valverde *et al*, 2019). Changes in glycosylation play a vital role in diseases such as increased fucosylation in prostate cancers (Höti *et al*, 2018; Saldova *et al*, 2011), dysregulated glycoforms in influenza virus (Roberts *et al*, 1993; Wan *et al*, 2019), varied glycosites of spike glycoprotein in COVID-19 (Wang *et al*, 2021; Watanabe *et al*, 2020), upregulated sialylation in cardiovascular disease (Gopaul & Crook, 2006), and elevated O-GlcNAcylation in neurodegenerative disease (Elbatrawy *et al*, 2020; Hart *et al*, 2011). In particular, protein glycosylation changes during tumorigenesis and cancer progression (Hakomori, 2002; Hakomori, 1989). Therefore, disease-specific glycosylation is often used as a diagnostic and/or prognostic biomarker. For instance, the core fucosylation of α-fetoprotein (AFP) is a clinical molecule for liver cancer diagnosis; using AFP core fucosylation instead of total AFP can improve sensitivity and specificity (Yin *et al*, 2014). Since most tumor markers approved by Food and Drug Administration (FDA) are glycoproteins, such as cancer antigen 125 (CA 125), AFP, immunoglobulins, neuron-specific enolase (NSE), and prostate specific antigen (PSA), potential cancer biomarkers are likely to be glycoproteins in human biofluids (Shiiki *et al*, 2011; Xu *et al*, 2021; Yang & Wang, 2017). Glycoenzymes (glycosyltransferases and glycosidases) may be intrinsically regulated in the tumor microenvironments (Chugh *et al*, 2015; Costa *et al*, 2020). Dysregulated glycoenzymes and their protein expression can alter protein glycosylation, leading to changes in the function of the protein cascade in the cell. Thus, analysis of tumor-specific glycosylation and upstream glycoenzymes is important to identify potential biomarkers for diagnosis and prognosis.

Glycoproteins are very abundant in biofluids, because when proteins are secreted from cells, they are likely to be post-translationally modified by glycans catalyzed by cellular glycoenzymes. Therefore, human plasma, urine, and saliva are used to discover disease-specific biomarkers. Tumor markers can be proteins or other substances that are present in or produced by cancer or other cells of the body in response to tumor microenvironment. Plasma markers such as PSA, CA-125, AFP or Amyloid-beta precursor protein (APP) have been clinically used for early detection of prostate cancer, ovarian cancer, liver cancers, and Alzheimer’s disease (German *et al*, 2007; Meany *et al*, 2009). Circulating tumors or tissue-specific tumors can also be used to evaluate markers for prognosis, diagnosis, or stages. Studies have shown that by detecting circulating tumor DNA methylation in a longitudinal study in patient’s plasma, early diagnosis of different cancers could be achieved (Chen *et al*, 2020). Recent studies have found that the expression of serum proteins (CEA (Carcinoembryonic antigen), RBP (Retinol-binding protein), and α1 antitrypsin) in the diagnosis of lung cancer has a sensitivity of 89.3% and a specificity of 84.7% (Patz *et al*, 2007). The results are based on analysis of the serum proteins of several patients diagnosed with non-small-cell lung cancer (NSCLC). However, more clinical studies are needed to confirm whether these results are applicable to different subtypes of NSCLC.

In addition to serum or plasma, which is widely used for biomarker discovery, saliva has become one of the essential biofluids in diagnosis due to non-invasive sample preparation. It can avoid the pain, anxiety or risk of infection, and it is easy to store and collect multiple subsequent specimens. Saliva has been used to diagnose oral diseases and monitor disease progression, such as periodontal pathogen (Kinney *et al*, 2011) or patients suspected COVID-19 (Fakheran *et al*, 2020; Yuan *et al*, 2020). Proteomic analysis of human saliva found that 48 out of the 500 proteins were differentially expressed between healthy controls (HC) and gastric cancer patients. Among them, STAT2 (signal transducer and activator of transcription 2) was up-regulated, and the tumor suppressor of DMBT1 (deleted in malignant brain tumors 1 protein) was down-regulated (Xiao *et al*, 2016). STAT family members such as STAT2 play an important role in the regulation of cell proliferation, differentiation, apoptosis and angiogenesis (Verhoeven *et al*, 2020). For example, upregulation of TLR2 driven by STAT3 can promote gastric tumorigenesis, and inhibition of STAT3 signaling can prevent gastric cancer proliferation and metastasis (Tye *et al*, 2012; Zhang *et al*, 2019b). A meta-analysis of 29 articles from more than 10,000 subjects showed that the diagnostic accuracy of saliva biomarkers for lung cancer remote from the mouth is up to 88% (Rapado-González *et al*, 2020). Therefore, saliva is a promising non-invasive biofluid for discovering novel biomarkers for lung cancer.

In addition to urea, ammonia, and electrolytes, saliva also contains many proteins. The most abundant saliva proteins are mucins, amylases, defensins, cystatins, histatins, proline-rich proteins, statherin, lactoperoxidase, lysozyme, lactoferrin, and immunoglobulins. These proteins can come from the salivary gland, stomach and lung (Wang *et al*, 2017; Xiao *et al*, 2012b). Mass spectrometry (MS) analysis of exosomes and macrovesicles in the saliva of lung cancer patients revealed that approximately 4% of the identified proteins belonged to distal lung cells. Among them, BPIFA1 (BPI fold-containing family A member 1), CRNN (Cornulin), MUC5B (Mucin-5B) and IQGAP (Ras GTPase-activating-like protein) are dysregulated in lung cancer, and most of which are also glycosylated (Sun *et al*, 2018). The changes in glycosylation may be attributed to the differential expression of glycoenzymes and their substrates in the tumor environment. Glycosyltransferases (GTFs), such as glucosyltransferase B (GtfB) (Smith *et al*, 2007), α1,3-fucosyltransferase (FUT5) (Gonzalez-Begne *et al*, 2011), α1,3-mannosyltransferase (ALG3), N-acetylgalactosaminide α2,6-sialyltransferase 1 (ST6GALNAC1), and α-N-acetyl-neuraminide α2,8-sialyltranserase 2 or 5 (ST8SIA2 or ST8SIA5) (the Human Protein Atlas), are highly abundant in saliva. Glycosylation of saliva-containing microbe, phagocyte, mucin or agglutinin is regulated by these GTFs (Cross & Ruhl, 2018). Saliva glycoproteins, MUC5B, MUC7 (mucin-7) (Tenovuo & Levine, 1989), salivary agglutinin (SAG) (Madsen *et al*, 2010), β-2-micoglobulin (Gussow *et al*, 1987), and proline-rich glycoprotein (PRG) (Tenovuo & Levine, 1989), can change when tumor initializes and progresses further through dysregulated glycoenzymes. Consequently, the identification of tumor-specific glycosylation and its dependent regulators is crucial for the discovery of biomarkers of interest.

Elucidating the glycosylation of saliva from healthy controls, non-cancer, and lung adenocarcinomas (ADC) patients is the key to revealing tumor-specific biomarkers. To decipher protein glycosylation, structural analysis of glycans, glycosites, site occupancy, and occupied glycans of glycosites is required. Glycan analysis can be performed by glycosidases or alkaline β-elimination (Jensen *et al*, 2012; Yang *et al*, 2017a), while N-glycosites are determined by tandem MS (MS/MS) against the intact N-glycopeptides enriched by hydrophilic interaction liquid chromatography (HILIC) (Riley *et al*, 2019; Xiao *et al*, 2018; Zacharias *et al*, 2016). Complex O-glycosylation has been successfully studied by O-protease, which cleaves the N-terminus of O-glycosylated serine or threonine; O-glycopeptides are usually analyzed by EThcD (Electron-transfer and higher-energy collision dissociation) fragmentation (Malaker *et al*, 2019; Yang *et al*, 2018; Yang *et al*, 2020). Conversely, the linkages of labile sialic acids are differentially derivatized by ethyl esterification and reductive amination using amine-containing compounds (Reiding *et al*, 2014; Yang *et al*, 2017b). The derivatization of sialic acid on the solid phase not only stabilizes the α-2,3 and α-2,6 linkages sequentially, but also facilitates the removal of reagents after the reaction (Yang *et al*., 2017b). By combining these analytical platforms and advanced MS technology, we can extensively deconvolute disease-specific glycopatterns by comparing protein glycosylation between healthy controls, non-cancer and cancer patients.

In this study, we used a solid-phase chemoenzymatic method to compare saliva glycosylation in healthy controls, non-cancers and cancer patients. To determine the linkage of fucosylation, glycoproteins are conjugated to a solid support and their fucoses are sequentially digested by specific α-fucosidases. Unstable sialic acids are modified by two-step chemical derivatization, and the linkages between α2,6 and α2,3 are distinguished by carrying a distinct mass tag after derivatization. Fucosylated glycoproteins are studied by bottom-up proteomics and matrix-assisted laser desorption/ionization (MALDI)-MS. Fucosyltransferases are quantitatively analyzed by qPCR. The biosynthesis of fucosylated high-mannose or complex N-glycans and their potential application for diagnosis of lung cancer are also discussed.

## Results

### Protein glycosylation differs between cancerous and non-cancer saliva

To show whether the glycosylation in the saliva of lung cancer patients has changed, we performed SDS-PAGE on the saliva proteins of healthy control (HC), other diseases (non-cancer, OD), and lung cancer (adenocarcinomas, LC) with and without glycosidase treatment. PNGase F (NEB BioLabs) is an N-glycosidase that can cleave all N-glycans from glycoproteins, but cannot cleave the innermost GlcNAc N-glycans with α1-3 fucose residue (e.g., plant or insect glycoproteins) (Tretter *et al*, 1993). SDS-PAGE showed that HC has more protein bands between 66 and 95 kDa, OD has fewer intensity bands between 66 and 70 kDa, and LC has high intensity proteins between 52 and 66 kDa (**Figure 3**). The protein pattern between 52 and 30 kDa is also different. After PNGase F digestion, the protein bands of all samples shifted to lower MW, indicating the presence of N-glycosylation in HC, OD and LC.

**Figure 1.**
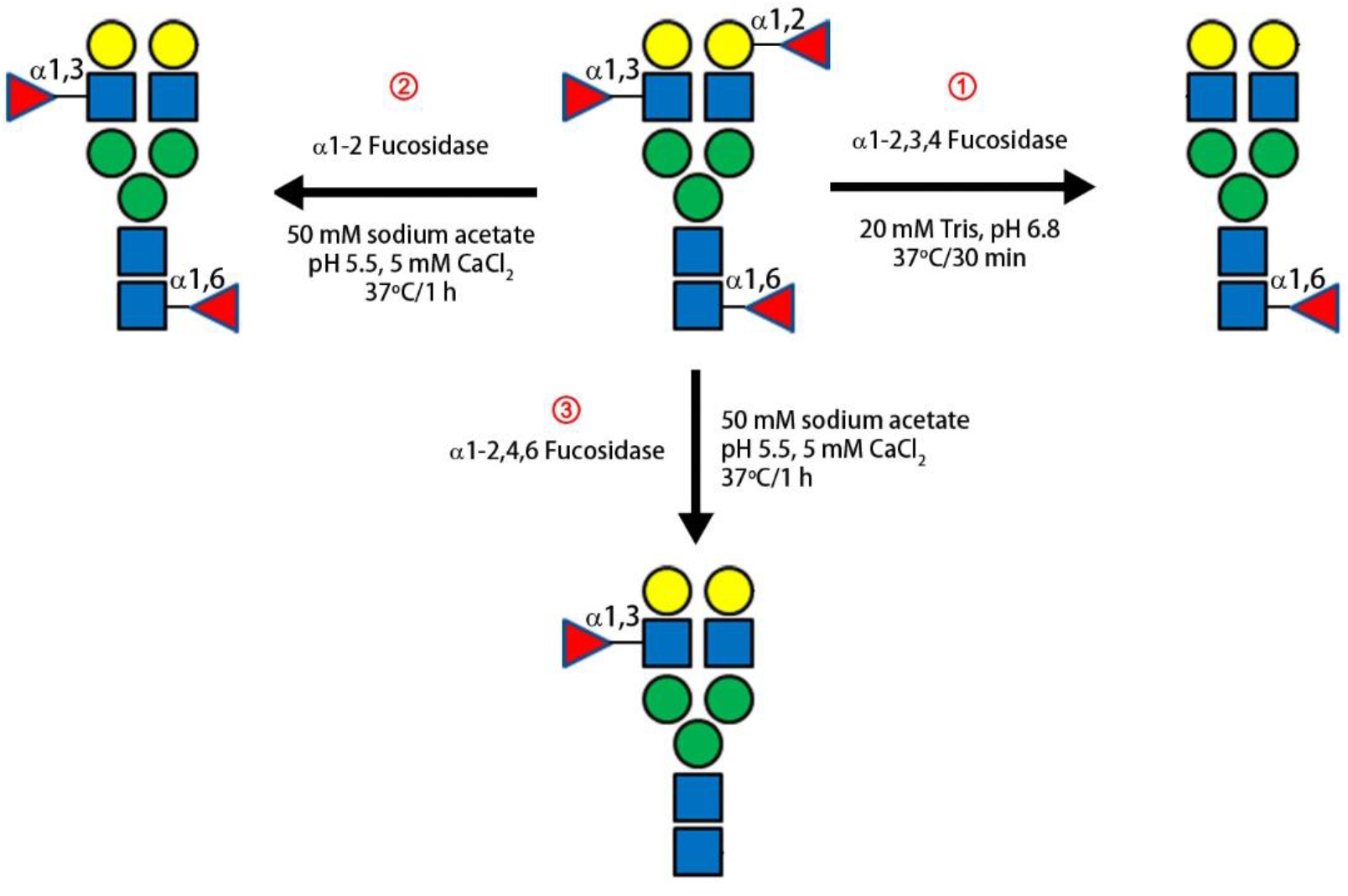
Schematic diagram of determination of fucosylation linkage using specific fucosidase. **①** Removal of all fucose linkages except for core α1,6 fucosylation by α1-2,3,4 fucosidase. This scheme led to the determination of core α1,6 linkage of fucosylated glycan; **②** Removal of α1,2 linkage of fucosylated glycan by α1-2 fucosidase. The remaining linkages of fucosylation can be α1,3 or α1,6. The α1,3 is then determined by comparing fucosylated glycans with scheme 1; **③** Removal of all linkages except for α1,3Fuc-GlcNAc through α1-2,4,6 fucosidase. This scheme confirms whether there is α1,4 linkage.

**Figure 2.**
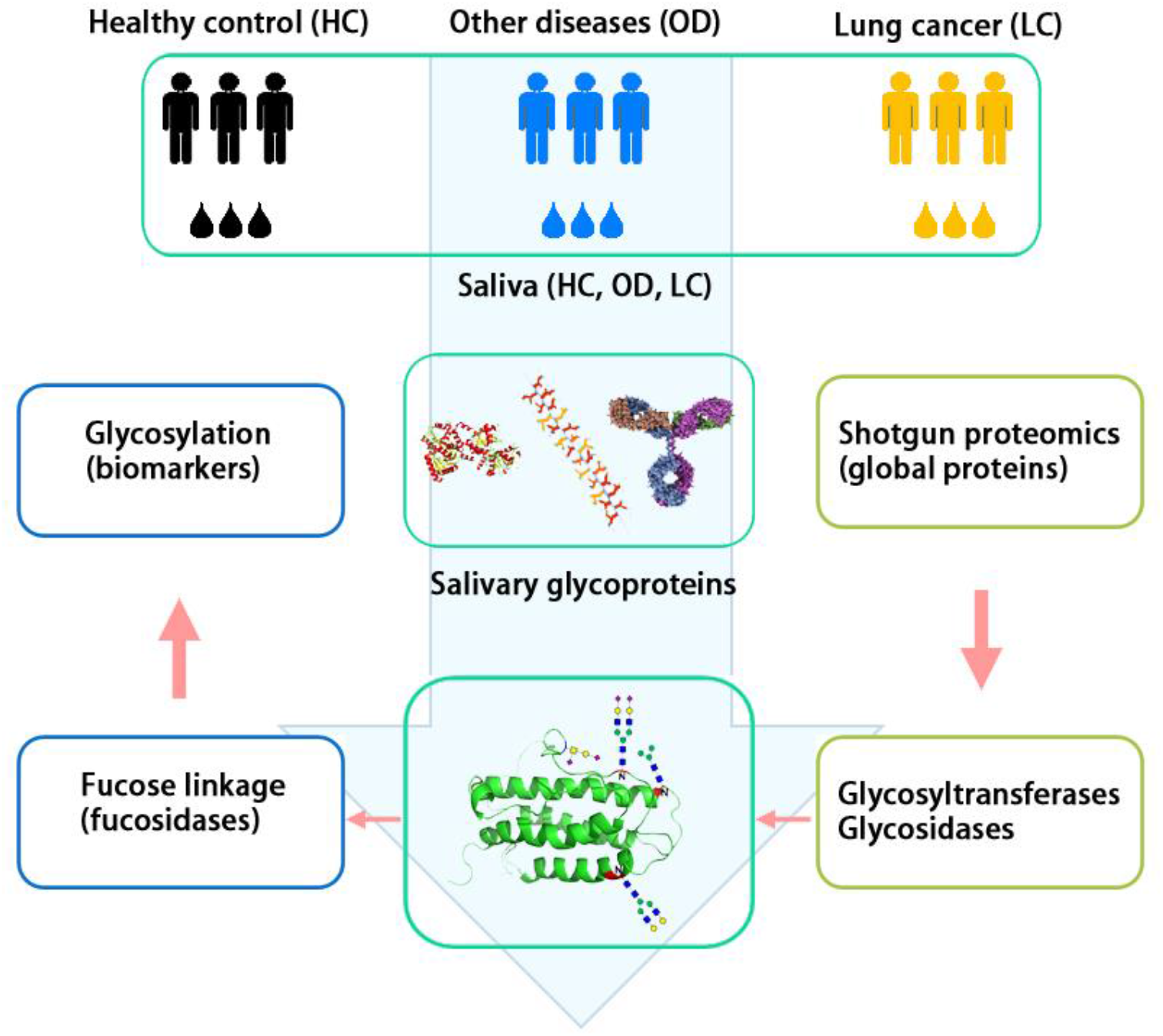
Workflow of mass spectrometric analysis of saliva proteins, glycoproteins and glycans. Three groups have been used for comparison, including healthy control (HC), other diseases (OD), and lung cancer (adenocarcinomas) (LC). Firstly, proteins were extracted from saliva and used for glycosylation analysis, bottom-up (or shotgun) proteomics and fucosylation linkage determination. Shotgun proteomics can identify glycosyltransferases responsible for specific glycosylation.

**Figure 3.**
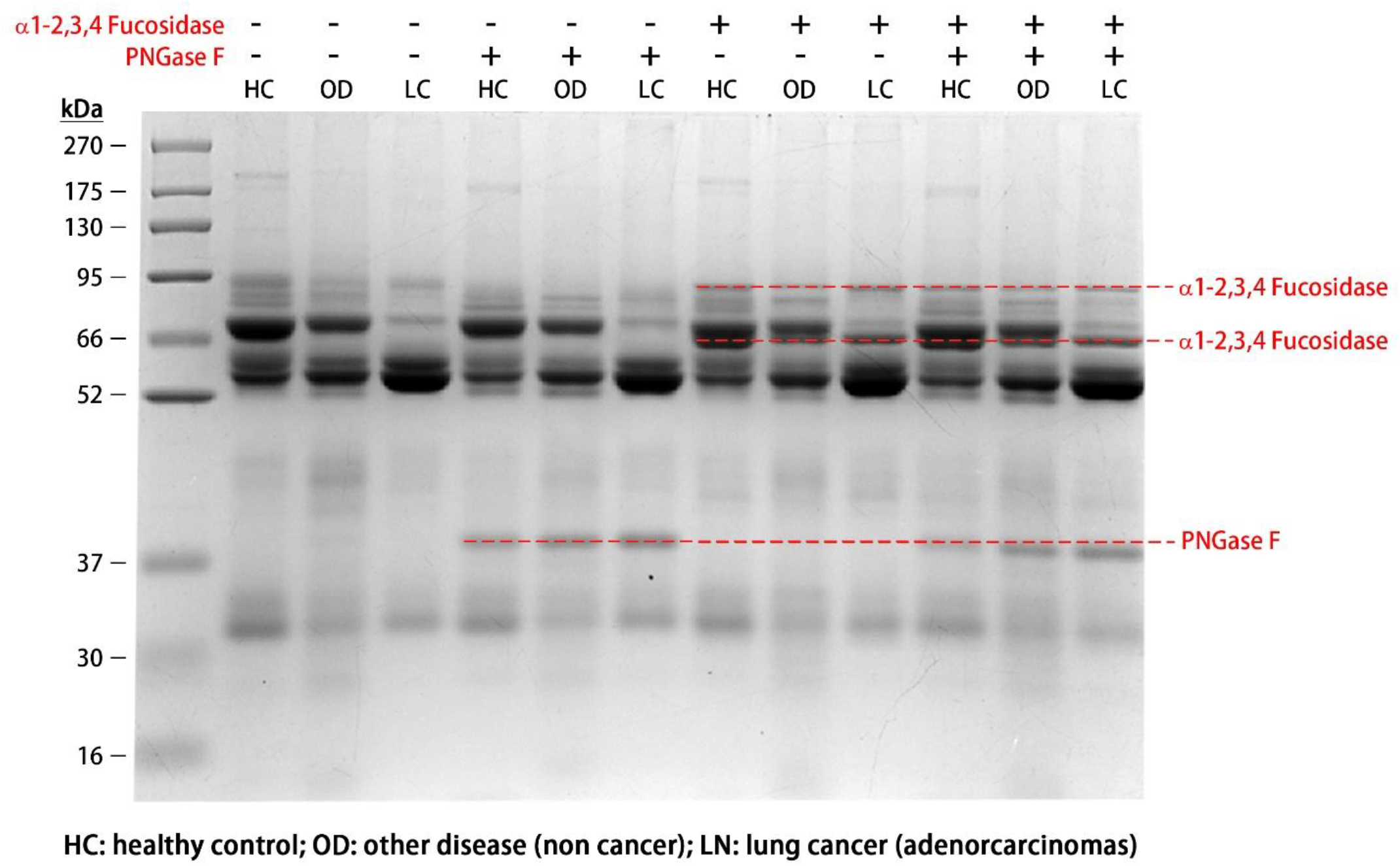
Different glycosylation present in saliva glycoproteins in healthy control, non-cancer, and lung cancer. Proteins of healthy control (HC), other disease (OD) and lung cancer (LC) were treated with PNGase F, α1-2,3,4 Fucosidase. PNGase F removes N-glycans from glycoproteins, thereby reducing the molecular weight (MW) of N-glycoproteins. α1-2,3,4 Fucosidase hydrolyzes fucose with linkages of α1-2, α1-3, or α1-4. A decrease in the MW of glycoproteins suggests one or more of these fucosylated linkages, but not α1-6. The MW of PNGase F is about 36 kDa, while α1-2,3,4 Fucosidase consists of two fucosidases modified with His-tags, with the MW of 87 kDa and 64 kDa (shown in the figure).

Fucosidase treatment can detect the presence of fucosylation on N-glycans or O-glycans, while fucosidase after PNGase F digestion can reveal whether there are fucoses on N-glycans. There was only a slight change when only fucosidase was used, but after treatment with both glycosidases, more pronounced protein bands appeared in OD and LC (**Figure 3**). These results indicate that fucosylation mainly occurs on N-glycosylation of lung cancer, and O-glycosylation containing fucosylation is negligible. The specific linkage of fucosylation can be further determined by fucosidase.

### Different linkages of fucosylated N-glycans are elevated in lung cancer

To determine the linkage of fucosylation, we used three fucosidases to process the glycoproteins on the solid support (resin) before PNGase F digestion (**Figure 1**). Because α1-2 fucosidase removes α1-2 fucose, the remaining linkage can be α1-3, α1-4 or α1-6. Similarly, α1-3 fucosidase can determine linkage α1-3 and the remaining linkage can be α1-2, α1-4 or α1-6. After α1-2,3,4 fucosidase digestion, any remaining fucose can be α1-6 core fucose. We used this strategy to elucidate the linkage of fucosylated N-glycans of saliva glycoproteins.

The glycan abundance after fucosidase treatment includes one that already exists in the sample and the other after corresponding fucose is removed. To explain our strategy, we used glycans with the same core structure H5N4 for quantitative analysis. As shown in **Figure 4**, the fucosylation linkage was determined by examining H5N4F4, H5N4F3, H5N4F2, and H5N4F1. The glycan profile of saliva glycoproteins indicates that these glycans are present in lung cancer, so removing any fucose will alter the relative abundance of the related glycans. For example, α1-2 fucosidase (F2) digests H5N4F4 to H5N4F2 (corresponding to two α1-2 linkages), or H5N4F3 to H5N4F2 and H5N4F1 (**Figure 4a**). Similarly, α1-2,3,4 fucosidase (F234) alters H5N4F4 to H5N4F3, H5N4F2, H5N4F1 (**Figure 4b**), resulting in the overall abundance of the final glycan profile (**Figure 4c**). The reduction of fucosylated glycans after F2 or F234 treatment indicates the presence of one or more of these fucosylation linkages in the sample. The core-fucosylated glycans were also identified because none of these fucosidases can digest α1-6 core fucosylation.

**Figure 4.**
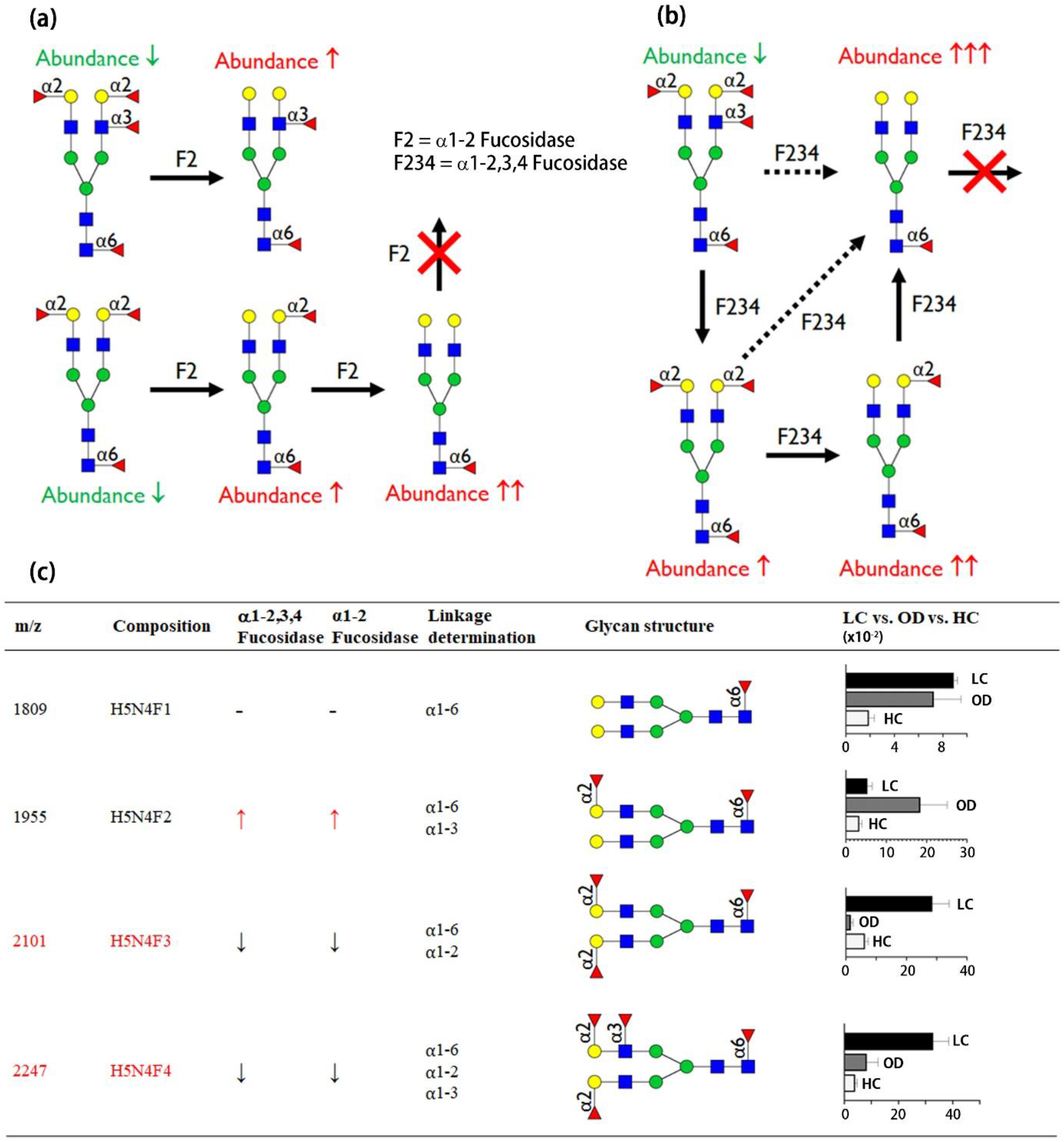
Determination of linkage of fucosylated glycans containing core structure of H5N4 using fucosidases. Saliva glycoproteins are extracted with lysis buffer and conjugated to the Aminolink plus resin. (**a**) Four fucosylated N-glycans (H5N4F4) contain α1-2 fucose on Gal after α1-2 fucosidase digestion, and the remaining fucose is either core α1-6 or antenna GlcNAc α1-3 linkage. F2 digestion can increase the abundance of H5N4F2, while H5N4F4 is reduced due to the loss of α1-2 fucose to form H5N4F2. Similarly, H5N4F3 loses two α1-2 fucose on Gal, and its abundance decreases accordingly. α1-2 fucosidase digestion eventually forms H5N4F1, while core α1-6 fucose still exists after F2 digestion. (**b**) α1-2,3,4 fucosidase (F234) determines the core-fucosylated glycan after removing α1-2, α1-3, and α1-4. H5N4F4 is trimmed to N5N4F3, N5H4F2, and H5N4F1. The abundance of these fucosylated glycans changes and is characterized by MALDI-MS. The remaining fucose of the glycan is a core fucose linkage. (**c**) The change of each glycan, H5N4F4, H5N4F3, H5N4F2, and H5N4F1 is the sum of these glycans and fucosidase digestion products in saliva. The reduction of H5N4F4 by F2 or F234 shows the presence of α1-2, α1-3, α1-4. The ratio of LC vs. OD. HC was a value measured from saliva glycoproteins without any fucosidase treatment. The arrow ↑ denotes the increase in abundance after fucosidase treatment compared with the untreated sample, and the arrow ↓ indicates decrease in abundance.

### Elevated fucosylation is unique to lung cancer

To confirm whether the glycan profile can be used to distinguish between LC and OD/HC, we analyzed the overall profile of glycans of the saliva glycoproteins from patients (**Supplementary Table S1**). The sample is processed on a solid support, and its sialic acids are stabilized by the ethyl esterification of α2,6-sialic acid and p-Toluidine carbodiimide coupling of α2,3-sialic acid (Yang *et al*., 2017b). The N-glycans are characterized by MALDI-TOF/TOF-MS and analyzed by GlycoWorkBench (Ceroni *et al*, 2008). **Figure 5** shows the glycan profiles of saliva glycoproteins in healthy controls (HC), lung cancer (LC) and other diseases (OD). Several conclusions can be drawn: 1) Compared with LC, HC has a lower glycan abundance because same amount of protein is conjugated to the resin. The highest peak observed in HC is H3N5F1 (1668 Da), and most glycans in LC are significantly higher than those in HC; 2) Most glycans are core-fucosylated, that is, α1,6 fucosylation to the innermost GlcNAc. These glycans have significantly higher intensity, such as H3N3F1, H3N4F1and H8N7F1; 3) Compared with HC or OD, the bisecting glycans with multiple fucosylation are obviously abundant in LC. These glycans include H4N3F3, H4N5F2, H4N5F3, H5N5F3, and H5N5F4; 4) There are multiple fucoses in core-GlcNAc (α1,6), antenna-GlcNAc (α1,3), and antenna-Gal (α1-2); 5) The glycan profile of OD is also different from that of HC. For example, the highest peak in HC is H3N5F1, and in OD it is H4N5F3. Generally, there are several fucosylation of glycans in OD compared to HC. These results indicate that the characteristics of fucosylated glycans can be used as markers to detect whether a patient has lung cancer or other diseases.

**Figure 5.**
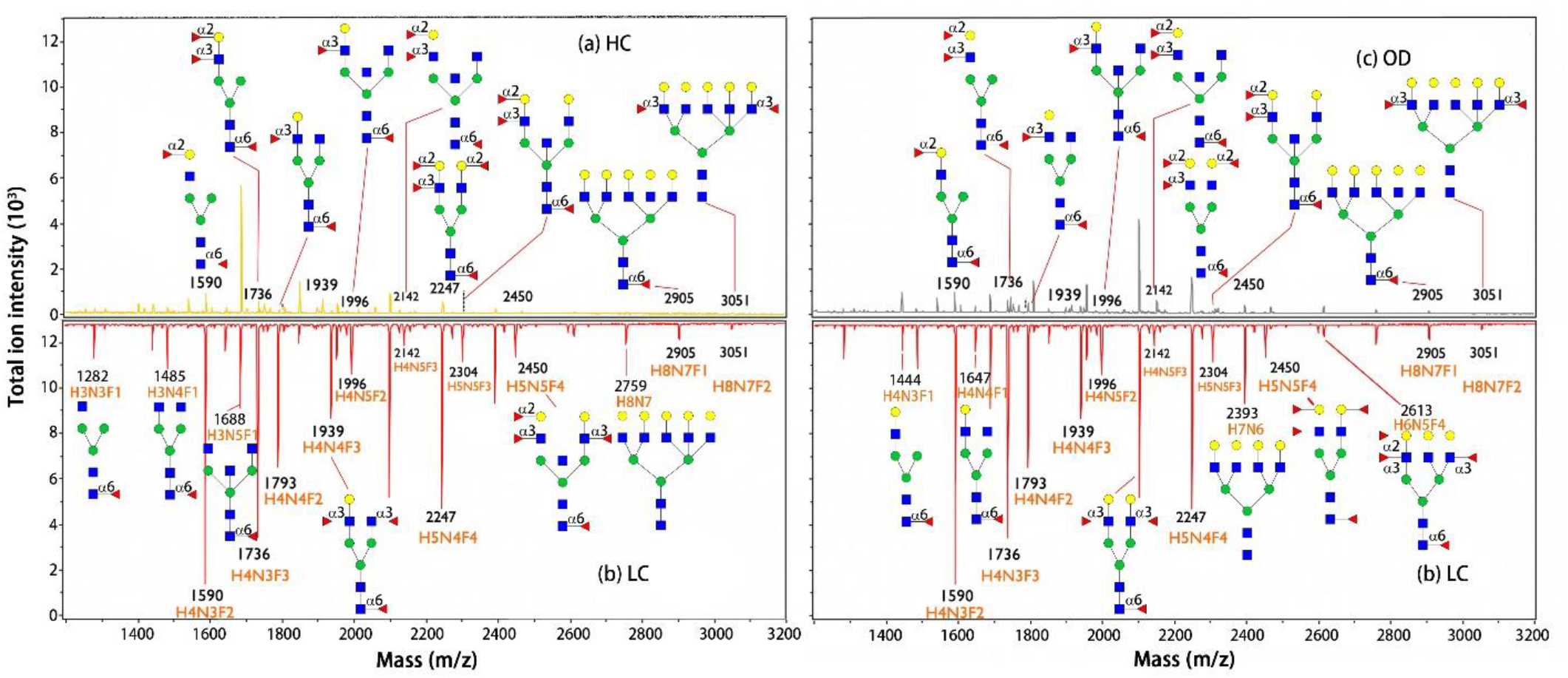
Different MALDI-MS glycan profile of saliva glycoproteins in cancer compared with the healthy control and other disease. (**a**) Glycan from healthy controls (HC). The glycan abundance of protein in HC is significantly lower than that of lung cancer (LC). (**b**) Glycan profile of saliva glycoproteins of lung cancer. The abundance of glycans with multiple fucosylation is significantly higher in cancer patients. These glycans also have core fucosylation, fucose on Gal or antenna GlcNAc. (**c**) Glycan profile of other disease (OD). The profile of OD is very different from that of LC. The characteristics of glycans can differentiate whether the patient is LC or OD.

The quantitative analysis of fucosylated glycans in HC, OD and LC is shown in **Table 1**, including glycan type, core structure of its fucosylated glycan, mass (MW), fucosidase digestion (F234 and F2), glycan structure, and abundance of fucosylated glycan of saliva without fucosidase treatment. We found that fucosylation occurs in high-mannose, hybrid and complex glycans. The core-fucosylation of Man3 (H3N2F1) decreased slightly in LC, but the changes in H4N2F1 and H5N2F1 were negligible. The core-fucosylated high-mannose (H3N2F1) was found in human saliva (Sinevici *et al*, 2019), and their possible biosynthetic pathways may involve FUT8 and α-mannosidase I (Nanno *et al*, 2020). Compared with those in HC, the OD of fucosylated high-mannoses is significantly reduced. Therefore, understanding the biosynthetic pathway of core-fucosylated high-mannose may be helpful for the diagnosis of non-cancer diseases using saliva.

**Table 1.**
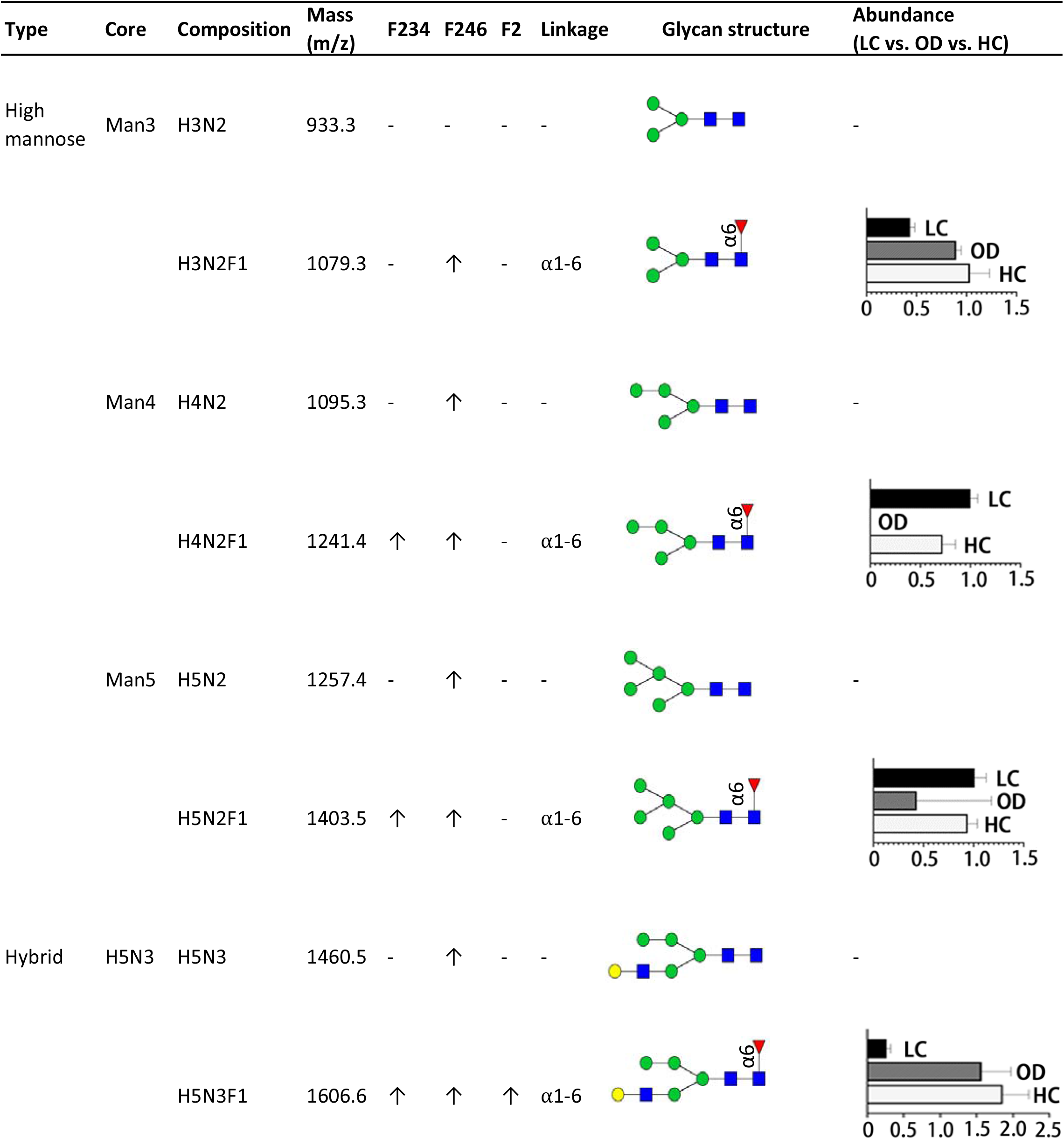
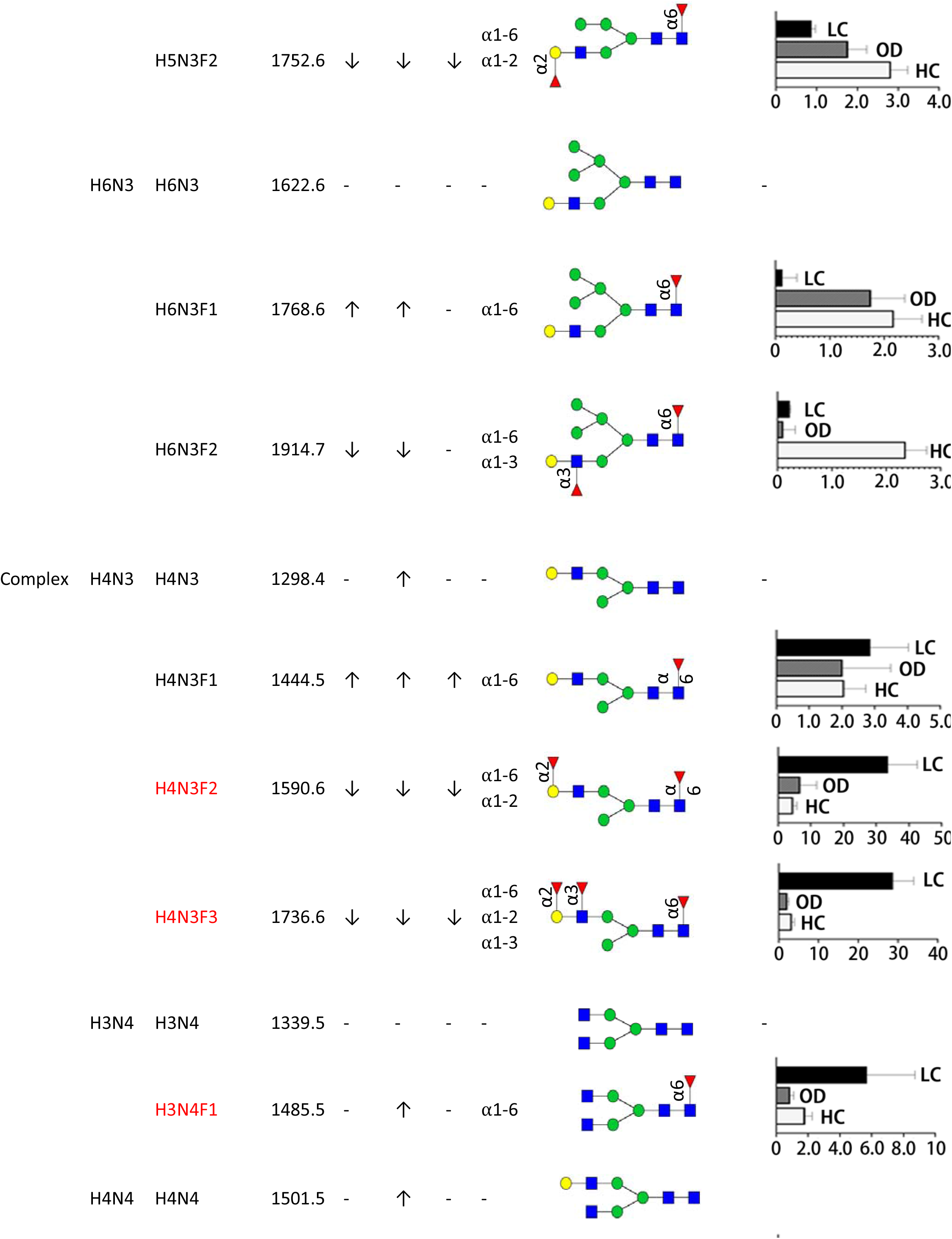
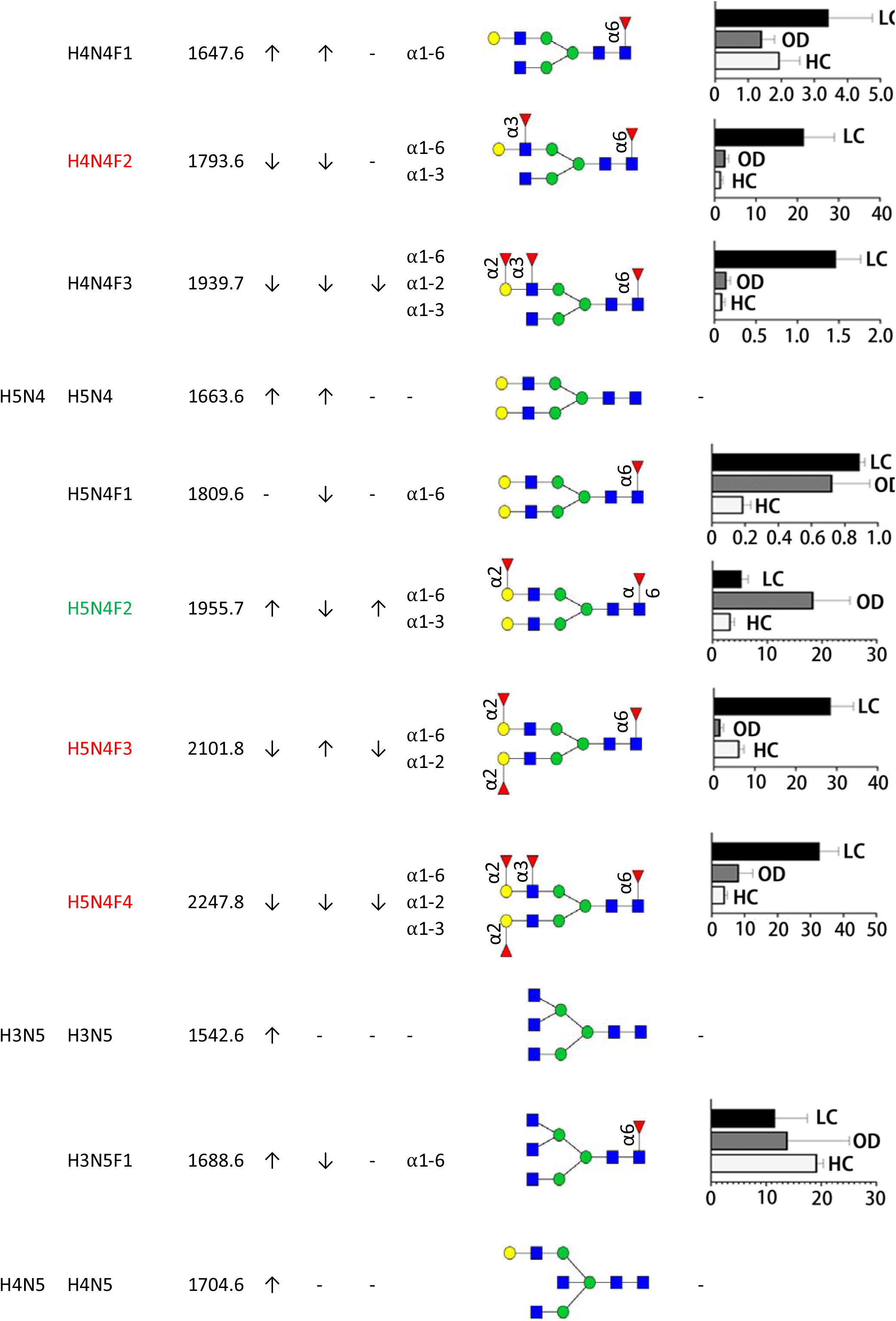
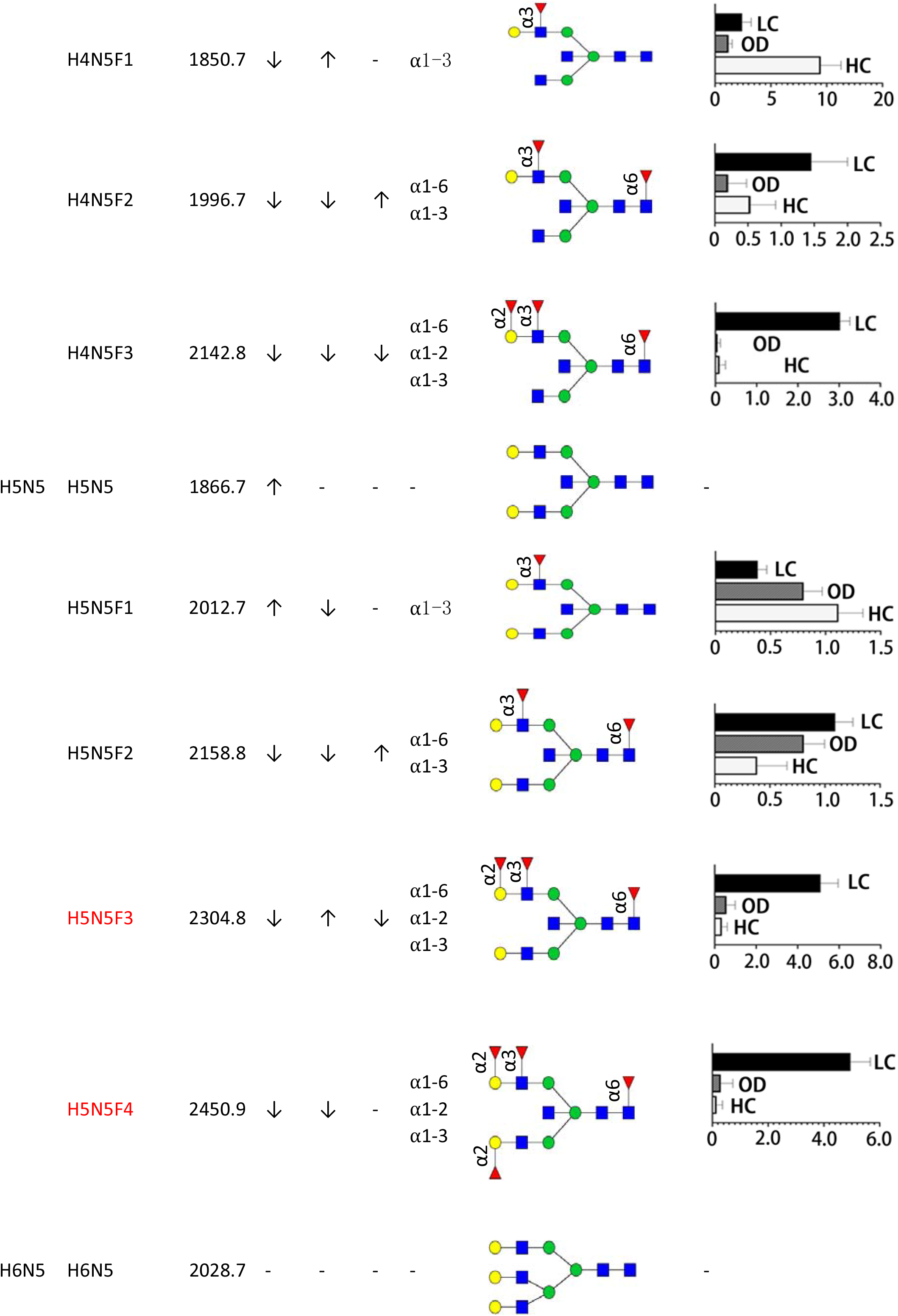
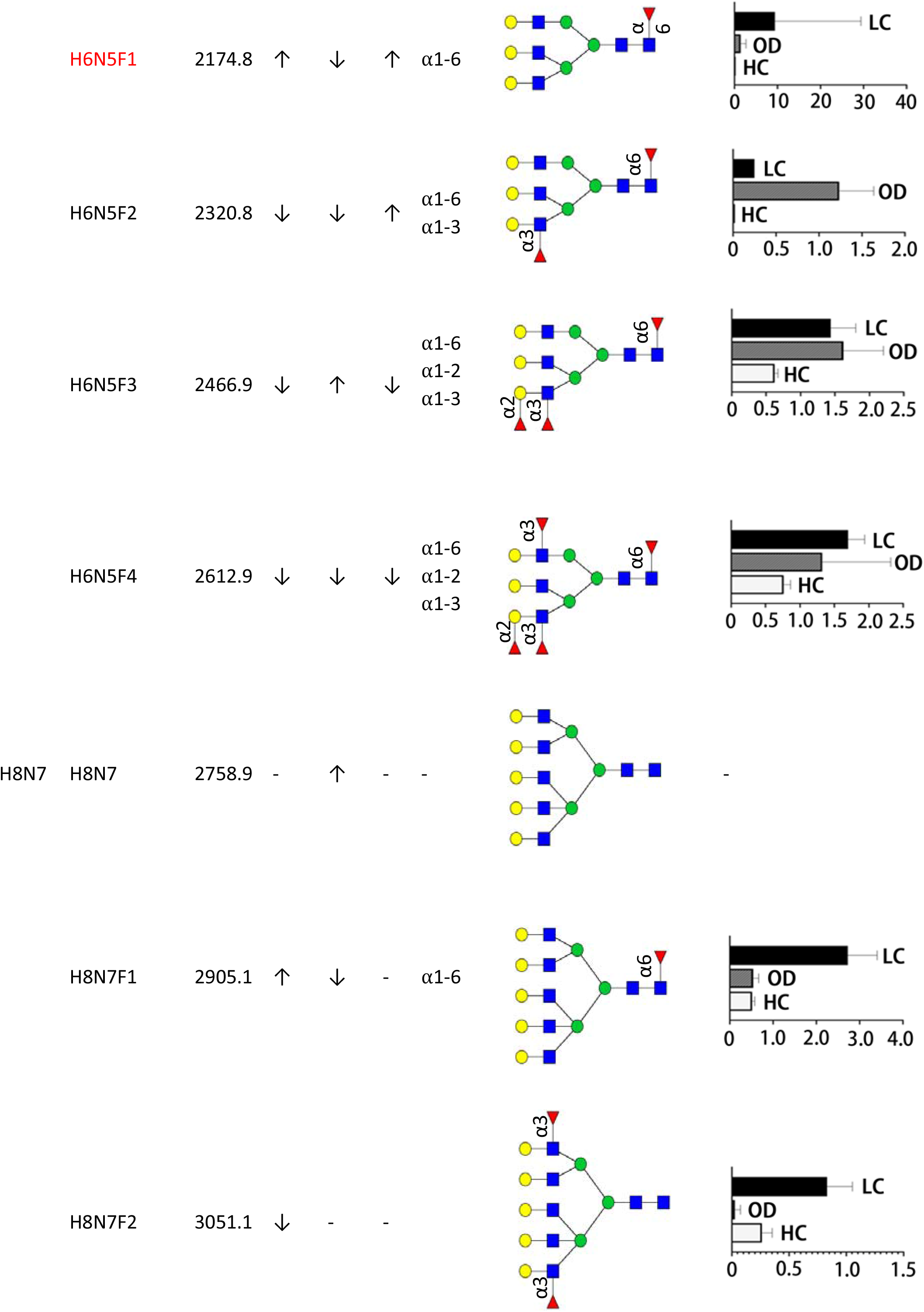
Regulation of fucosylated glycans of saliva glycoproteins in lung cancer compared with other diseases and healthy controls. The linkage of fucose was determined by fucosidase. The abundance between LC, OD and HC was measured by MALDI−MS without any fucosidase treatment. The measurement was conducted in triplicate. H = Hex, N = HexNAc, F = Fucose, F234 = α1−2,3,4 Fucosidase, F246 = a1−2,4,6 Fucosidase, F2 = α1−2 Fucosidase. The arrow ↑ and ↓ stand for increase or decrease of glycan after fucosidase treatment (10^−2^).

Hybrid glycans were detected in human saliva, and they were greatly reduced in lung cancer (**Table 1**). The four fucosylated hybrid glycans H5N3F1, H5N3F2, H6N3F1, and H6N3F2 have much lower intensity in LC saliva. These glycans feature a core fucose, two of which have α1-2 fucose on Gal or α1-3 on GlcNAc. A similar trend was observed in gastric cancer serum, where the hybrid glycan H6N4F1 was down-regulated in cancer serum (Ozcan *et al*, 2014). How these hybrid glycans are regulated in tumorigenesis and cancer progression remains to be discovered, but they lack N-acetylglucosaminyltransferase I (GnT1), an enzyme responsible for the synthesis of hybrid and complex glycans, which can lead to delayed embryonic development (Ioffe & Stanley, 1994).

The most striking changes were observed in complex glycans with at least one fucose. We list 25 complex glycans that are significantly upregulated in lung cancer (**Table 1**). Except for H3N5F1, H4N5F1 and H5N5F1, most of the glycans in LC are higher than those in OD or HC. The dominant increase in complex glycans is those with two or more fucoses, such as H4N3F2, H4N3F3, H4N4F2, H5N4F3, H5N4F4, H5N5F3, and H5N5F4. Because these complex glycans have core fucose, this suggests that the core fucosylation enzyme FUT8 is actively regulated in cancer. Studies have shown that the expression of FUT8 in tumor lesion is upregulated in NSCLC and is associated with tumor metastasis or malignancy (Chen *et al*, 2013). According to the Human Protein Atlas, FUT8 protein is highly abundant in lung and digestive tract tissues, and its mRNA is highly expressed in salivary gland, tongue and lung; however, proteomic analysis of saliva proteins did not detect FUT8 in HC, OD or LC. These results suggest that the core-fucosylated proteins should come from the lungs or other organs. Additionally, the formation of α1,2-linked fucose on Gal or the formation of α1,3-linked fucose on GlcNAc was observed in saliva glycoproteins of lung cancer patients. The increase in fucosylation specific for LC should be attributed to the increase in the expression of the corresponding fucosyltransferases (FUTs), which have been further characterized by qPCR.

### Upregulated fucosyltransferases lead to aberrant fucosylation in lung cancer

To identify FUTs, we used clinical specimens form lung tumor tissue and matched adjacent non-tumor tissues to quantify the mRNA level of each FUT. To determine whether the saliva contains FUTs for synthesizing linkages of fucose, we used shotgun proteomics to analyze FUT expression in HC and LC. LC-MS/MS data showed that FUT6 and FUT11 were present in the saliva proteins of LC patients, but no other FUTs were identified from saliva (**Supplementary Table S3** and **Table S4**). In contrast, the abundance of mRNA extracted from saliva is extremely low. As a result, we did not observe any FUT mRNA expression using saliva samples.

Proteomic analysis of human saliva shows that there is an inherent correlation between the protein components of lung tissue and saliva. Literature studies have shown that when people suffer from lung cancer, protein signature appears in human saliva (Xiao *et al*, 2012a). The presence of specific glycosylation can be traced back to lung tissue. To this end, we use qPCR to quantitatively characterize FUTs in lung tissue. As shown in **Figure 6a**, 8 FUTs were found in adjacent non-tumor tissue (non-cancer) and tumor tissue (lung cancer). Among these FUT genes, FUT4, FUT6, FUT7 and FUT9 are highly expressed in lung cancer, while FUT1 and FUT3 have limited increase. Interestingly, the change in FUT8 mRNA levels between LC and HC is negligible, although the core-fucosylation in LC is significantly higher than in HC.

**Figure 6.**
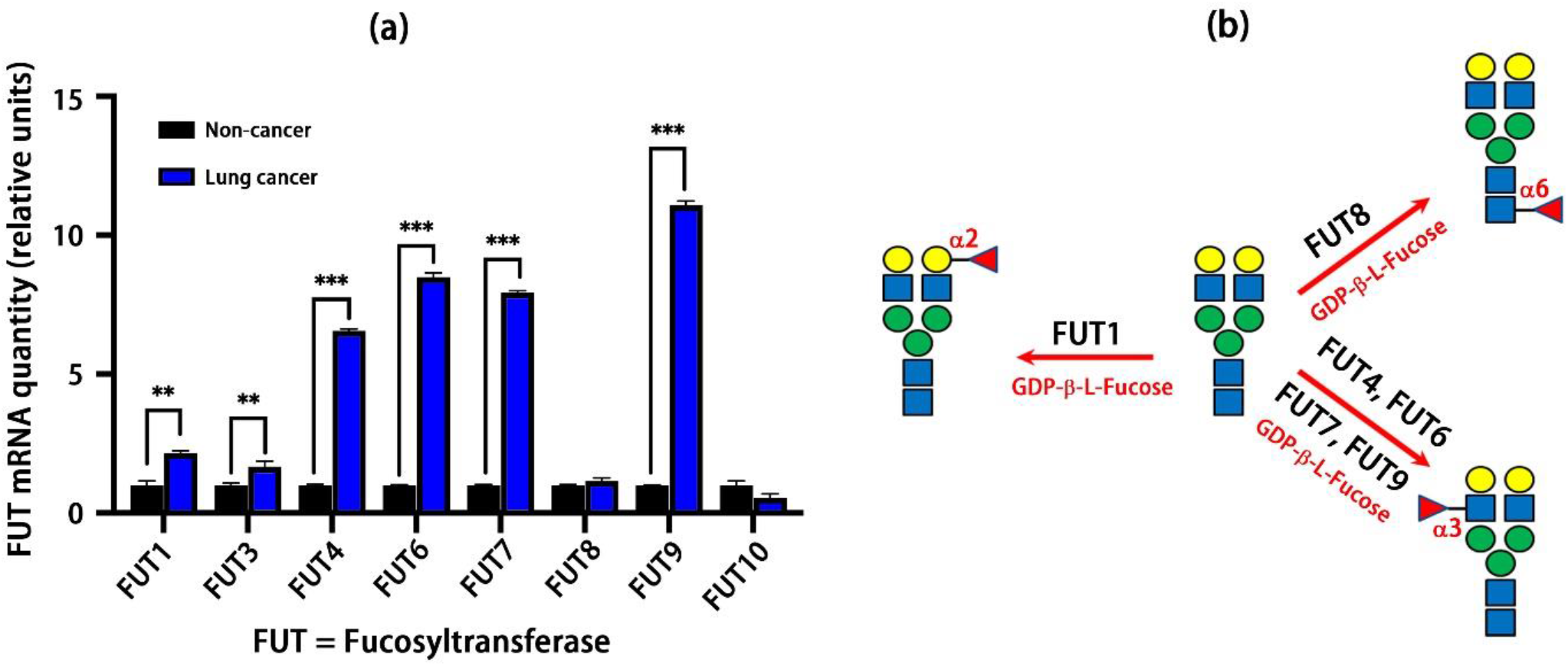
The biosynthetic pathway of formation of different fucosylation linkage by fucosyltransferase (FUT) enzymes through qPCR quantification. (**a**) The qPCR quantification of FUT genes showed a substantial increase in FUT1 (2.17-fold), FUT3 (1.68), FUT4 (6.56), FUT6 (8.49), FUT7 (7.93), FUT9 (11.09). The mRNA is extracted from lung tissues of adjacent non-tumors (control) and tumors; (**b**) FUT8 enzyme transfers GDP-β-L-Fucose to the innermost GlcNAc, forming core-fucosylation (1.16-fold). The α1,2 linked fucosylation is catalyzed by FUT2, and the α1,3 linked fucosylation is catalyzed by the combination of FUT4, FUT6 and FUT9 enzymes.

The different linkages of fucosylation are regulated by specific FUT enzymes. **Figure 6b** schematically shows the biosynthetic pathway for fucosylation via various FUTs. Theoretically, FUT1 or FUT2 catalyzes GDP-β-L-Fucose to Gal, forming α1,2-linked fucosylation (Goupille *et al*, 2000; Tan *et al*, 2016). mRNA expression indicates that FUT1 is an enzyme that synthesizes α1,2-linked fucosylation in lung tissue. FUT8 is responsible for the synthesis of α1,6-linked fucosylation and exists in LC and HC. FUT8 is associated with unfavorable clinical outcomes and may be a prognostic marker of lung cancer (Honma *et al*, 2015). FUT4, FUT6 and FUT9 are the three main isoforms that catalyze the α1,3-linked fucosylation in lung cancer. Some N-glycans (**Figure 5**) have α1,2, α1,3 and α1,6-linked fucosylation, indicating that these FUTs are highly expressed in lung cancer.

## Discussion

Our study shows that aberrant fucosylation is manifested in saliva glycoproteins of lung cancer. The characteristics of fucosylated glycans are quite different from those of healthy controls or other diseases. Most glycans have increased core and antenna fucosylation in lung cancer. Although many studies have reported the upregulation of α2,6-linked sialic acids in lung cancer serum (Vasseur *et al*, 2012; Zhang *et al*, 2018), the increase in sialylation of lung cancer saliva glycoproteins is negligible. The saliva glycoproteins, such as Mucin-5B, IgA, lactotransferrin, zinic-a2-glycoprotein etc.(Cross & Ruhl, 2018), do possess sialic acid residues, but in our research, we found that dominant change is fucosylation. Our data shows that the characteristics of fucosylation can distinguish whether a patient has lung cancer or other diseases (**Figure 5**).

Because the tumor microenvironment alters the expression of glycoenzymes, abnormal fucosylation has been reported in various cancers. Importantly, fucosylation plays a vital role in cancer biology by regulating tumor signal transduction and cell-cell adhesion pathways, and performs tumor immune surveillance through necrosis factor-related apoptosis-inducing ligand signaling (Zhang *et al*, 2019a). Fucosylation analysis of prostate cancer cell lines showed that FUT1 is highly elevated compared to normal prostate cells and is regulated in LNCaP, so glycans carrying α1,3-linked fucose are elevated in prostate cancer (Fukushima *et al*, 2009). Changes in the expression of fucosyltransferases (FUTs), FUT1, FUT3, FUT6 and FUT8, are associated with poor diagnosis and tumor metastasis in NSCLC (Park *et al*, 2020). Therefore, it is significant to identify FUTs in saliva and how this altered expression affects fucosylation.

The fucosylation is formed by transferring a GDP-β-L-Fucose to the substrate catalyzed by a specific fucosyltransferase. As shown in **Figure 6b**, three different fucose linkages are catalyzed by the respective enzymes. It is known that FUT8 can synthesize α1,6 Fuc-GlcNAc which is the core-fucosylated N-glycans. However, more than one FUT enzyme can catalyze the transfer of GDP-β-L-fucose to Gal or antenna GlcNAc. For instance, FUT1 is responsible for the synthesis of α1,2 Fuc-Gal, any one of FUT4, FUT6, FUT7 and FUT9 can synthesize α1,3 Fuc-GlcNAc. Studies have shown that knocking down the FUT1 gene can attenuate tumor cell proliferation in HER2-overexpressed NCI-N87 cells (Kawai *et al*, 2013). Similarly, the upregulation of FUT1 in lung cancer may lead to an increase in α1,2 fucosylation in lung cancer saliva. In summary, our study shows that 1) FUT8 in lung cancer leads to an increase in the level of core fucosylation, 2) FUT1 upregulation is the main driving factor for the significant increase in α1,2 linkage fucosylation, 3) FUT4, FUT6, FUT7 and FUT9 are highly upregulated to elevate expression of α1,3 linked fucosylation.

Due to the unique characteristics of fucosylation in lung cancer, the different glycan profiles between lung cancer and healthy control/other disease can be used for the diagnosis of lung cancer. Since each fucosylated glycoform can be recognized by a different lectin, a microarray or lectin-based ELISA (enzyme-linked immunosorbent assay) can be used to quantify and determine fucosylated linkage. Our future work includes the use of lectins, such as Lens Culinaris Agglutinin (LCA) (α1,6), Ulex Europaeus Agglutinin I (UEAI) (α1,2), or Aleuria Aurantia Lectin (AAL) (α1,2, α1,3, α1,4, α1,6), to study linkage-specific glycoproteins. Additionally, collecting saliva from early patients may help determine the characteristics of fucosylation for early diagnosis.

## Conclusions

Our study shows that aberrant fucosylation of saliva glycoproteins defines lung cancer malignancy. Since the proteins in human biofluids are highly glycosylated, attempts are made to identify disease-specific markers through changes in protein glycosylation in biofluids. Abnormal glycosylation is usually produced by dysregulated glycoenzymes, which are responsible for adding or removing monosaccharides to or from glycans. The tumor microenvironment can cause glycoenzyme dysregulation that is very different from the normal pathophysiological state. Lung cancer tends to have higher FUT expression, leading to up-regulation of fucosylation. Glycoproteomics and glycomic analysis of saliva indicate that aberrant fucosylation is unique to lung cancer, while other diseases (such as lung inflammatory) or healthy controls show a distinct fucosylation than lung cancer. Our results confirmed that the increase in FUT1 expression enhanced α1,2-linked fucosylation, while FUT4,6,7,9 catalyzed the upregulation of α1,3-linked fucosylation. In contrast, FUT8 mRNA expression is also present in LC and adjacent non-tumor tissue, which indicates that FUT8 mRNA alone is not sufficient as a marker of lung cancer, rather than using fucosylation patterns for tumor diagnosis.

## Methods and Protocols

Unless specified otherwise, all materials were purchased from Aladdin (Shanghai, China). Glycoprotein standard and acetonitrile was purchase from Tedia (Fairfield, Ohio, USA). Trifluoroacetic acid (TFA) and chloroform were purchased from Lingfeng Chemical Reagent (Shanghai, China). Anhydrous ethanol and hydrochloric acid were purchased from Qiangsheng (Suzhou, Jiangsu, China). Methanol was purchased from Hushi (Shanghai, China). Fetuin, α1-2 fucosidase, α1-2,4,6 fucosidase, PNGase F and α2-3,6,8 neuraminidase was purchased from New England BioLabs (Ipswich, MA, USA). α1-2,3,4 fucosidase (FucosEXO) was purchased from ACROBiosystems Inc (Shanghai, China). RevertAid First Strand cDNA Synthesis Kit was purchased from Thermo Scientific and SYBR qPCR Master Mix was purchased from Vazyme (Nanjing, Jiangsu, China). Human GAPDH endogenous reference genes primers were purchased from Sangon Biotech (Shanghai, China). All primers used for qPCR were synthesized by GENE WIZ Company (Suzhou, Jiangsu, China). 10% SDS-PAGE Gel SuperQuick preparation kit, pre-stained color protein ladder, SDS-PAGE sample loading buffer, bicinchoninic acid (BCA) protein assay kit, TRIzol, diethyl pyrocarbonate (DEPC) water and 1M Tris-HCl were purchased from Beyotime (Haimen, Jiangsu, China).

### Saliva protein extraction

All patient samples were collected according to the protocols approved by the Institutional Review Board (IRB) of Soochow University and the written informed consent was provided to patients in advance. The sample was processed and stored in a similar manner. The 500 μL solution consists of TCA (trichloroacetic acid; 20% w/v), acetone (90% v/v), and DTT (dithiothreitol; 20 mM), and was mixed with 500 μl saliva. The mixture was vortexed and precipitated overnight at −20°C. The sample was then centrifugated at 15,000 rpm for 30 min at 4°C. The supernatant was discarded and the pellet was collected, then washed with 200 μL of cold acetone (90%) and 20 mM DTT, and finally washed with cold acetone (80%) and 10 mM DTT. To suspend the pellet in the solution, the sample was sonicated for at least 5 min prior to acetone-DTT wash. The pellet was placed at −20°C for 20 min, then centrifugated at 15,000 rpm for 5 min at 4°C. Finally, the pellet was collected and dried in Speed-Vac (5 min) and stored at −80°C before further analysis.

### SDS-PAGE and glycosidase treatment of saliva proteins

The concentration of saliva proteins was measured by BCA assay and nanodrop. The three groups of saliva samples were diluted to a concentration of ~1mg/ul. 20 μg proteins were taken from each group and reacted with PNGase F, fucosidase and the mixture of the two enzymes at 37°C for 4 h. The untreated saliva and the enzymically digested saliva sample were mixed with 5X protein loading buffer and incubated at 100°C for 5 min. Electrophoresis was performed on SDS-PAGE using 10% sample and 10% SDS-Page Gel Kit (Beyotime). The running buffer is composed of 0.025 M Tris, 0.192 M Glycine, and 0.1% SDS. 20 μl of sample and loading buffer mixture was added to each well. After electrophoresis, the gel was stained in the staining solution (containing 0.25% Coomassie Bright Blue R250, 45% methanol and 10% acetic acid) for 3 h, and then eluted in the eluting buffer (methanol:glacial acetic acid:water = 2:2:9, V/V) until the protein band was clear. Then the gel imager (Bio-Rad) was used for gel band imaging.

### Enzymatic release of glycans

Glycans are released from glycoproteins through chemical derivatization and enzymatic digestion on solid-phase (Yang *et al*., 2017a; Yang *et al*, 2013). Briefly, the protein (500 μg) is heated at 90°C for 10 min and then mixed with 200 μL of AminoLink plus resin, which is pre-conditioned with 500 μL of 1x binding buffer (twice). The 1x binding buffer contains 10 mM sodium citrate and 5 mM sodium carbonate. The protein is conjugated to the resin in 1x binding buffer (4 h at room temperature (RT)), followed by adding 50 mM NaCNBH_3_. Buffer is changed to 1x PBS and the resin is further incubated 4 h in 1x PBS in the presence of 50 mM NaCNBH_3_. The active sites of the resin were blocked with 1 M Tris.HCl (pH 7.4). Sialic acids are derivatized by 0.25 M EDC/0.25 M HBot in ethanol at 37°C/1h, then modified by 1 M p-Toluidine (pT) (Yang *et al*., 2017b). After washing, the resin is treated with glycosidases to analyze fucosylation linkages or glycan composition.

### Determination of fucosylation linkage

The fucosylation linkage of glycoproteins conjugated to the resin can be further determined by fucosidase and MS (**Figure 1**). The linkage is resolved by α1-2 fucosidase, α1-2,3,4 fucosidase or α1-2,4,6 fucosidase. The conjugated glycoprotein was aliquoted into three equal amounts and treated with three fucosidases. An aliquot was incubated in 50 unit of α1-2,3,4 fucosidase in 20 mM Tris.HCl (pH 6.8), 37°C/30min. N-glycans were released by 0.2 μL PNGase F in 200 μL of 20 mM NH_4_HCO_3_, 37°C/overnight (**Figure 1**①). The second aliquot was treated with 10 units of α1-2 fucosidase in 50 mM sodium acetate and 5 mM CaCl_2_ (pH 5.5), 37°C/1h, and then by PNGase F to release N-glycans (**Figure 1** ②). The third aliquot was treated with 10 units of α1-2,4,6 fucosidase under same condition and its N-glycans were released by PNGase F (**Figure 1**③). The linkage of α1,2, α1,3 and α1,6 is thus determined.

**Mass spectrometry analysis of glycans and glycoproteins**

After the PNGase F incubation, the glycans were eluted by centrifugation and further washed with 100 μL of HPLC water (twice). The total volume is approximately 400 μL, of which 2-4 μL is used for glycan analysis by Bruker AutoFlex MALDI-TOF/TOF Mass Spectrometry. Each sample is tested 3-4 technical duplicates, and each measurement performs an average of 10,000 shots. Global proteins are analyzed by shotgun proteomics. In short, the protein (500 μg) was dissolved in 8M urea and treated with 12 mM TCEP (Tris (2-carboxyethyl) phosphine hydrochloride) (37°C/1h), and 16 mM IAA (iodoacetamide) (RT/1h in the dark). 10 μg trypsin (Promega, Madison, WI, USA) was added to the protein after dilution (< 1.5 M urea). Trypsin digestion was conducted overnight at 37°C and peptides were further purified by C18 SPE (solid-phase extraction) column. Site-specific glycosylation analysis was performed by solid-phase extraction of glycopeptide enrichment (SPEG) (Zhang *et al*, 2003). The purified peptides were oxidized by 10 mM sodium periodate for coupling of glycopeptides to hydrazide beads. Glycan-containing glycopeptides were released by PNGase F. The peptides were analyzed by Thermo Scientific Orbitrap Fusion LC-MS, using the same parameters described in our previous work (Yang *et al*., 2018).

### Comparison of fucosylation of saliva glycoproteins

The workflow of clinical samples was shown in **Figure 2**. Saliva specimens include samples from healthy controls (HC), other non-cancer disease (OD), and lung adenocarcinomas (LC) (see **Supporting Information Table S1**). Proteins are extracted from saliva according to the Saliva Protein Extraction protocol. The proteins (1 mg) were used to determine the fucosylation linkage using a solid-phase chemoenzymatic method (**Figure 1**). The aliquot proteins (500 μg) were also digested with trypsin for quantitative analysis of glycosyltransferases. The structure of glycans in HC, OD and LC was compared for features that are specific to cancer.

### qPCR quantification of fucosyltransferases in lung tissue

The fucosyltransferases of interest were quantitatively analyzed by q-PCR using the ABI 7500 Real-Time PCR instrument. The TRIzol method was used to extract total RNA from lung cancer tissues and matched adjacent non-tumor tissues. RNA concentration was measured using Nanodrop. The extracted RNA was reversed into cDNA using the RevertAid First Strand cDNA Synthesis Kit (Thermo). The primer sequences for qPCR are shown in the **Supplementary Table S2**. We use human GAPDH as the reference gene. The reaction system is 10 μl 2× ChamQ Universal SYBR qPCR Master Mix, 0.4 μl of 10 μM upstream and downstream primers, 1 μl of cDNA template, and 20 μl of water for the final system. The reaction procedure of the qPCR system is as follows: pre-deformation at 95°C for 30 s; 40 cycles of amplification (95°C for 10s, 60°C for 30 s); melting curve (60°C for 60s, 95°C for 15 s). After the reaction, relative gene expression was calculated quantitatively by 2^^-^ΔΔ^Ct^.

## Acknowledgements

This work was supported by the startup funding of Soochow University, Jiangsu Province-Suzhou Science and Technology Planning Project SL T201917. We thank the Priority Academic Program Development of the Jiangsu Higher Education Institutes (PAPD).

## Authors’ contributions

Z. Gao: Formal analysis, investigation, visualization. Z. W: MALDI-MS analysis, visualization. Y. Han: LC-MS/MS, visualization. X. Zhang: Resources, supervision, methodology, review and editing. P. Hao: Resources, supervision, methodology, review and editing. M.M. Xu: Investigation, visualization, supervision, software. S. Huang: Investigation, visualization. S.W. Li: Investigation, visualization, review and editing. J. Xia: resource, visualization, review. J.H. Jiang: Conceptualization, resources, funding acquisition, formal analysis, review and editing. S. Yang: Conceptualization, resources, funding acquisition, formal analysis, methodology, writing-original draft, writing-review and editing.

